# Apolipoprotein ε4 modifies obesity-related atrophy in the hippocampal formation of cognitively healthy adults

**DOI:** 10.1101/2021.11.12.468385

**Authors:** Bethany M. Coad, Parisa A. Ghomroudi, Rebecca Sims, John P. Aggleton, Seralynne D. Vann, Claudia Metzler-Baddeley

## Abstract

Characterizing age- and risk-related hippocampal vulnerabilities may inform about the neural underpinnings of cognitive decline. We studied the impact of three risk-factors, Apolipoprotein (*APOE)*-ε4, a family history of dementia, and central obesity, on CA1, CA2/3, dentate gyrus (DG) and subiculum in 158 cognitively healthy adults (38-71 years). Subfields were labelled with the Automatic Segmentation of Hippocampal Subfields (ASHS) and FreeSurfer (version 6) protocols. Volumetric and microstructural measurements from quantitative magnetization transfer and Neurite Orientation Density and Dispersion Imaging were extracted for each subfield and reduced to three principal components capturing apparent myelin/neurite packing, size/complexity, and metabolism. Aging was associated with an inverse U-shaped curve on myelin/neurite packing and affected all subfields. Obesity led to reductions in myelin/neurite packing and size/complexity regardless of *APOE* and FH status. However, amongst individuals with a healthy Waist-Hip-Ratio, *APOE* ε4 carriers showed lower size/complexity than non-carriers. Protocol type did not affect this risk pattern. These findings provide novel evidence for interactive effects between *APOE* and central obesity on the hippocampal formation of cognitively healthy adults.

**Highlights:**

- Age-related inverted U-shaped curve of hippocampal myelin/neurite packing
- Obesity-related reductions of hippocampal myelin/neurite packing and size/complexity
- *APOE* modifies the effects of obesity on hippocampal size/complexity
- Age-related slowing of spatial navigation
- No *APOE*, family history, or obesity effects on cognition

## 1. Introduction

The world’s population is aging, creating an increase in age-related health issues, including cognitive decline (Beard et al., 2016). With respect to age-related memory impairments, the hippocampal formation, i.e., dentate gyrus, cornu Ammonis (CA) fields, and subiculum, warrants particular attention as it is critically involved in memory processing and is affected early in the progression of Alzheimer’s disease (AD). Magnetic resonance imaging (MRI) studies have, for example, shown reductions in total hippocampal volume associated with aging in cognitively healthy individuals (Raz, 2001), while hippocampal atrophy remains one of the supporting diagnostic features of amnestic Mild Cognitive Impairment (aMCI) and AD (de Flores et al., 2015a). The next challenge is to distinguish normal age-related hippocampal changes from potential pathological changes related to genetic and lifestyle risk factors of dementia in pre-symptomatic individuals (Jack et al., 2013). Overall structural volumes may, however, lack sufficient sensitivity for early detection. Instead, multi-parametric quantitative MRI indices may provide an alternate route to detect subtle microstructural changes that, in turn, aid our understanding of both age- and dementia risk-related effects on the hippocampal formation and its subfields (Kodiweera et al., 2016; Metzler-Baddeley et al., 2019a; Metzler-Baddeley et al., 2019b; Nazeri et al., 2015; Wolf et al., 2015).

The objectives of this study were three-fold: Firstly, to characterize the pattern of age and age-independent dementia risk effects on hippocampal subfield macro- and microstructure in a sample of cognitively healthy adults (38-71 years of age) (Metzler-Baddeley et al., 2019a; Metzler-Baddeley et al., 2019b). The effects of three established risk factors of dementia, i.e., carriage of the Apolipoprotein E (*APOE)* ε4 genotype (Angelopoulou et al., 2021; Chai et al., 2021; Feringa and van der Kant, 2021; Koutsodendris et al., 2021; Liu et al., 2013), a positive family history (FH) of dementia in a first-grade relative (Alosco et al., 2014; Donix et al., 2010; Johnson et al., 2014; Tanzi, 2012; Wolf, 2012), and abdominal obesity (Arnoldussen et al., 2014; Beydoun et al., 2008; Chuang et al., 2016; Pedditizi et al., 2016; Xu et al., 2011) and their potential interactions were studied (Metzler-Baddeley et al., 2019a; Metzler-Baddeley et al., 2019b; Mole et al., 2020).

Secondly, to investigate the pattern of age and risk effects not only with volumetric but also with multi-parametric microstructural MRI from diffusion neurite density and dispersion imaging (NODDI) (Zhang et al., 2012) and quantitative magnetization transfer (qMT) (Eng et al., 1991; Henkelman et al., 1993; Henkelman et al., 2001; Sled, 2017) to aid our understanding of the biophysical properties underpinning aging and risk (Wolf et al., 2015). We also assessed whether the choice of protocol for the labelling of hippocampal subfields had an impact on the analysis of risk effects (Yushkevich et al., 2015a). For this purpose, the main subfields of the hippocampal formation, i.e., CA1, CA2/3, dentate gyrus (DG) and subiculum were segmented with two publicly available, automated protocols that make use of T_1_- and T_2_-weighted hippocampal images: The Bayesian inference labeling methods implemented in FreeSurfer 6.0 (Iglesias et al., 2015) and the Automatic Segmentation of Hippocampal Subfields (ASHS) that utilizes multi-atlas segmentation and machine learning techniques (Yushkevich et al., 2015b). Both protocols have recently been validated with histopathological evidence in patients with epilepsy (Menzler et al., 2021; Mizutani et al., 2021).

Thirdly, to explore whether individual differences in hippocampal subfield macro- and microstructure correlated with episodic memory and spatial navigation abilities known to rely on hippocampal processes (Brown et al., 2014; Hartley et al., 2005; Hoang et al., 2018; Kyle et al., 2015).

The evidence regarding age-related atrophy in hippocampal subfields remains mixed. Volume reductions have previously been reported for CA1 (Malykhin et al., 2017; Mueller and Weiner, 2009; Raz et al., 2015; Uribe et al., 2018; Wisse et al., 2014), CA2-4 (Daugherty et al., 2016; Malykhin et al., 2017; Mueller and Weiner, 2009; Raz et al., 2015; Wisse et al., 2014), subiculum (de Flores et al., 2015b; La Joie et al., 2010; Malykhin et al., 2017; Wolf et al., 2015) and DG (de Flores et al., 2015b; La Joie et al., 2010) but were not consistently observed across all studies. A recent UK Biobank data analysis found non-linear age-related changes in all subfields of the hippocampal formation (Veldsman et al., 2021). In this study, female *APOE* ε4 homozygotes over the age of 65 years exhibited the largest atrophy across CA1, CA3, CA4, subiculum and presubiculum, suggesting that age and sex modulated the effects of *APOE* (Veldsman et al., 2021) (see also Donix et al., 2010; Dounavi et al., 2020; Kerchner et al., 2014; Mueller et al., 2008; Mueller and Weiner, 2009; Reiter et al., 2017). *APOE* ε4-related volume reductions in the molecular layer of the subiculum and the CA fields were also observed in a middle-aged cohort of cognitively healthy participants, while no effects were present for FH or cardiovascular risk (Dounavi et al., 2020) (Donix et al., 2010; Dounavi et al., 2020; Kerchner et al., 2014; Mueller et al., 2008; Mueller and Weiner, 2009; Reiter et al., 2017).

Furthermore, lifestyle-related factors, such as obesity and sedentary lifestyle, may have adverse effects on the hippocampus. For instance, abdominal visceral fat has been found to be associated with volume reductions (Anan et al., 2010) and increases in free water signal of the whole hippocampal formation (Metzler-Baddeley et al., 2019a). Obesity is related to pro-inflammatory states (Cox et al., 2015) and in rodent models was found to induce microglia activation and reduce long-term potentiation in the hippocampus (Hao et al., 2016). It is increasingly recognized that *APOE* ε4 may interact with obesity to augment disruptions in lipid, glucose, insulin, and immune response metabolism which in turn may increase AD risk (Jones and Rebeck, 2018; Mole et al., 2020; Zade et al., 2013). Consistent with this view, we have previously found interaction effects between *APOE*, FH, and obesity on white matter microstructure, that were particularly pronounced in the right parahippocampal cingulum (Mole et al., 2020). More specifically, *APOE* ε4 carriers with a positive FH exhibited obesity-related reductions in apparent myelin, while no effects were observed for those without a FH. These risk-effects were moderated by hypertension and inflammation-related blood markers. The parahippocampal cingulum is a limbic white matter pathway that connects the posterior cingulate and parietal cortices with the medial temporal lobes (Bubb et al., 2018) and is known to be affected in amnestic Mild Cognitive Impairment (Metzler-Baddeley et al., 2012; Yu et al., 2017). It is therefore possible that adverse interactions between genetic and lifestyle risk factors may increase the vulnerability of medial temporal lobe structures to neurodegeneration. However, the nature of these interactions and their impact on the subfields of the hippocampal formation remain poorly understood and require further elucidation.

To this purpose we studied MRI data from 158 asymptomatic individuals from the Cardiff Aging and Risk of Dementia Study (CARDS) (38-71 years of age) (Coad et al., 2020; Metzler-Baddeley et al., 2019a; Metzler-Baddeley et al., 2019b; Mole et al., 2020), a sample well-characterized with regards to their genetic and lifestyle risk of dementia (Mole et al., 2020). MRI measures consisted of intracranial volume (ICV) adjusted hippocampal subfield volumes and the following microstructural indices: i) NODDI intracellular signal fraction (ICSF) for apparent neurite density, ii) NODDI isotropic signal fraction (ISOSF) estimating free water, iii) NODDI neurite orientation dispersion index (ODI), iv) qMT macromolecular proton fraction (MPF) for apparent myelin (Ceckler et al., 1992; Schmierer et al., 2007), v) qMT forward exchange rate *k_f_* estimating tissue metabolism (Giulietti et al., 2012; Harrison et al., 2015) and vi) longitudinal relaxation rate R_1_ for apparent water, lipid/protein, and, to a lesser extent, iron content (Callaghan et al., 2015).

Support for studying neurite properties comes from neuropathological evidence suggesting that human aging is associated with a reduction of neocortical dendritic spine density (Dickstein et al., 2013) with accompanying compensatory increases in the dendritic extent of DG granular cells (Flood et al., 1985; Flood et al., 1987). Likewise, recent *in vivo* NODDI studies found age-related reductions in neocortical neurite dispersion (Nazeri et al., 2015) and increases in neurite dispersion of the whole hippocampus (Metzler-Baddeley et al., 2019b; Nazeri et al., 2015). With regards to AD pathology, ODI and ICSF were found sensitive to amyloid and tau pathology in the hippocampus (Colon-Perez et al., 2019) and in white matter in AD animal models (Colgan et al., 2016; Colon-Perez et al., 2019). In addition, qMT *k_f_* reductions were reported in the hippocampus, temporal lobes, parietal cortex and posterior cingulate in people with AD (Giulietti et al., 2012) and T_1_ and T_2_ relaxometry measurements have been found sensitive to AD pathology in animal and human imaging studies (Knight et al., 2016; Knight et al., 2019; Tang et al., 2018). In previous CARDS analyses, we observed age-related reductions in left hippocampus *k_f_* and R_1_ (Metzler-Baddeley et al., 2019b) as well as *APOE* ε4-related reductions in left thalamus qMT MPF (Mole et al., 2020). These observations demonstrate that microstructural MRI indices may provide complementary information to volumetric measurements capturing age and risk-related differences in metabolic, macromolecular, and free water-related tissue properties (Callaghan et al., 2015; Giulietti et al., 2012; Harrison et al., 2015).

It should be noted though that our previous CARDS analyses did not find risk effects on microstructural measurements of the whole hippocampal formation (Mole et al., 2020; see also Dounavi et al., 2020, Evans et al., 2020). An important limitation of these studies was the treatment of the hippocampal formation as a unitary structure, while this region comprises multiple subfields that are thought to be differentially vulnerable to age and disease-related processes. Notably the subiculum and CA1 regions have been proposed to be particularly vulnerable in aMCI and AD (Adler et al., 2018; Khan, et al., 2015; Lindberg et al., 2017) and in asymptomatic individuals with positive amyloid and tau cerebrospinal fluid (CSF) biomarkers (Tardif et al., 2018).

However, one source of inconsistency when subdividing the hippocampal formation stems from different subfield and border segmentation protocols (Yushkevich et al., 2015a). Yushkevich et al. (2015a) compared 21 protocols for labelling hippocampal subfields, including those adopted here (ASHS and FreeSurfer). They found considerable differences with regards to the region within each segmentation was performed, the set of the employed anatomical labels, and the extents of specific anatomical labels. The largest discrepancies between the protocols were at the CA1/subiculum boundary and in the anterior portion relative to body and tail portions of the hippocampal formation.

For this reason, we employed the two most widely used automated hippocampal subfield labelling protocols, i.e., FreeSurfer version 6.0 and ASHS. We then assessed whether the type of protocol affected the pattern of risk-related differences in macro- and microstructure of the hippocampal formation. We focused our comparison on CA1, CA2/3, DG and subiculum as these regions have previously been implicated in aging and disease and were sufficiently large to extract meaningful microstructural information from diffusion and qMT images with a resolution of approximately 2mm^3^.

While the microstructural measurements described above were chosen to capture complementary tissue properties (Wolf et al., 2015), they also share some overlapping information which can cause redundancies in the data analysis and reduce the statistical power of the analyses (Chamberland et al., 2019). We and others have previously demonstrated that Principal Component Analysis (PCA) can be successfully employed to reduce the dimensionality of multi-modal brain measurements and extract meaningful components of the underlying data structure (Bourbon-Teles et al., 2017; Chamberland et al., 2019; Geeraert et al., 2020; Metzler-Baddeley et al., 2017; Penke et al., 2010).

Here we adopted this approach to study potential dissociations between age- and age-independent risk effects on principal components of hippocampal macro- and microstructure. More specifically, we modelled main and interaction effects of *APOE*, FH, WHR, protocol and hippocampal subfields on three principal components that reflected apparent myelin/neurite packaging, size/complexity and metabolism of hippocampal gray matter. These analyses controlled for the effects of age, sex (Veldsman et al., 2021) and verbal intelligence (Boyle et al., 2021), as we aimed to gain a better understanding of age and sex-independent effects of *APOE*, FH and obesity and their potential interactions. Finally, we assessed age and risk effects on cognitive components of episodic memory and spatial navigation and explored brain-function correlations between the hippocampal subfield macro-and microstructure and cognition.

## 2. Materials and Methods

CARDS received ethical approval from the School of Psychology Research Ethics Committee at Cardiff University (EC.14.09.09.3843R2). Participants provided written informed consent in accordance with the Declaration of Helsinki.

### 2.1 Participants

Participants between the age of 38 and 71 years were recruited from the local community *via* Cardiff University volunteer panels, notice boards and local poster advertisements. A detailed description of the CARDS sample can be found in Mole et al. (2020). All participants had a good command of the English language and were without a history of neurological and/or psychiatric disease, head injury with loss of consciousness, or drug or alcohol dependency. A total of 166 CARDS volunteers underwent MRI scanning at the Cardiff University Brain Research Imaging Centre (CUBRIC). Seven participants did not complete the MRI protocol, including the high resolution T_2_ images of the hippocampus, due to claustrophobia and/or feeling uncomfortable. Participants’ intellectual function was assessed with the National Adult Reading Test - Revised (NART-R) (Nelson, 1991) and cognitive impairment was screened for with the Mini Mental State Exam (MMSE) (Folstein et al., 1975). One person with a MMSE score of 26 was excluded from the analysis. Thus, the current analysis was based on 158 datasets (see Table 1).

**Table 1.**
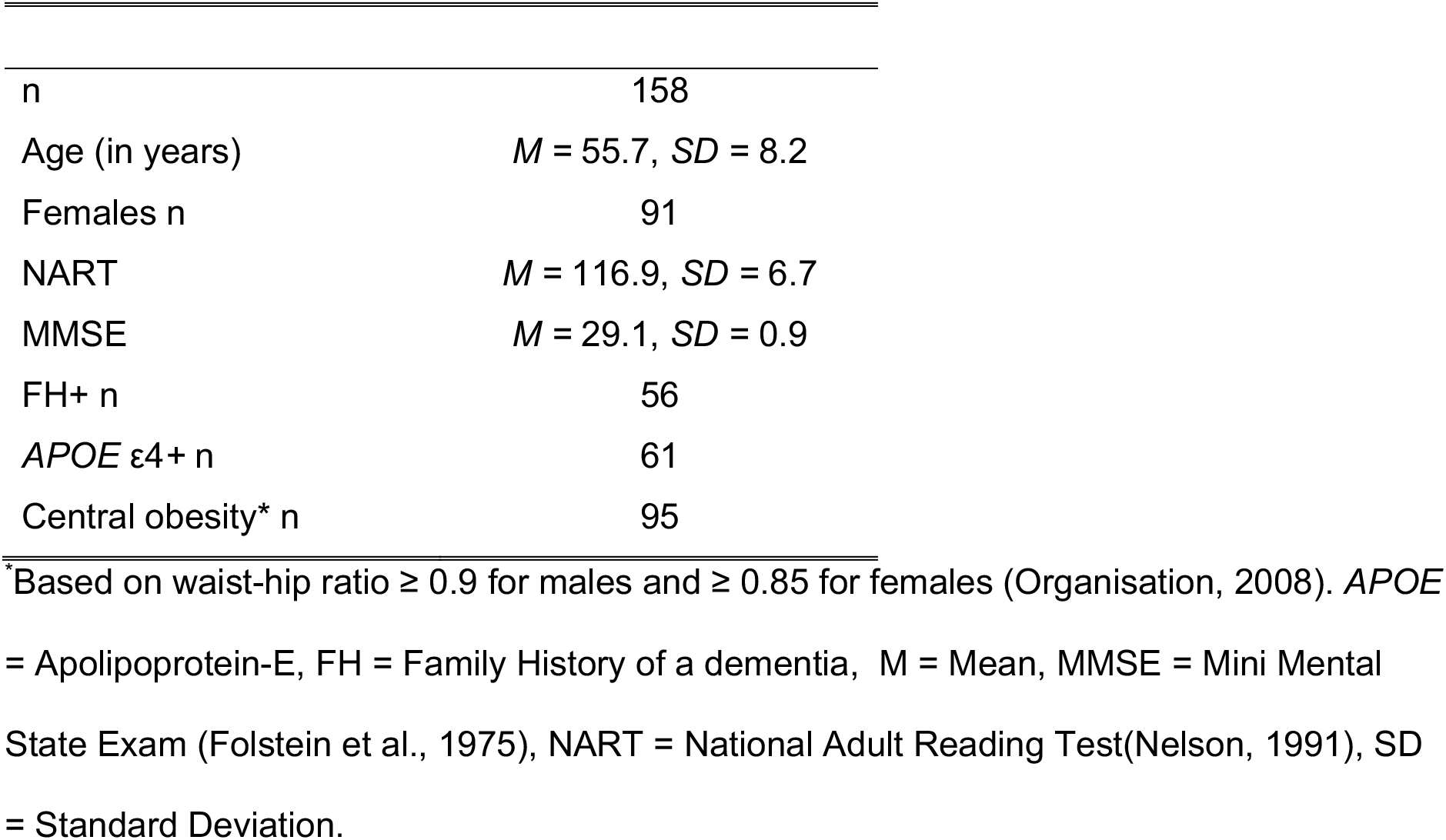
Summary of demographic, genetic, and lifestyle risk information of participants.

|  |  |
| --- | --- |
| n | 158 |
| Age (in years) | $M = 55.7, SD = 8.2$ |
| Females n | 91 |
| NART | $M = 116.9, SD = 6.7$ |
| MMSE | $M = 29.1, SD = 0.9$ |
| FH+ n | 56 |
| *APOE* $\epsilon 4$+ n | 61 |
| Central obesity* n | 95 |

\*Based on waist-hip ratio $\geq 0.9$ for males and $\geq 0.85$ for females (Organisation, 2008). *APOE* = Apolipoprotein-E, FH = Family History of a dementia, M = Mean, MMSE = Mini Mental State Exam (Folstein et al., 1975), NART = National Adult Reading Test (Nelson, 1991), SD = Standard Deviation.

### 2.2 Assessment of dementia risk factors

Central obesity was assessed with the Waist to Hip Ratio (WHR) following the World Health Organisation’s (Organisation, 2008) recommended protocol for measuring waist and hip circumference. Central obesity was defined as a WHR ≥ 0.9 for males and ≥ 0.85 for females. Individuals were categorized as centrally obese (WHR+) or normal WHR (WHR-).

Saliva samples were collected with the Genotek Oragene-DNA kit (OG-500) for DNA extraction and *APOE* genotyping. *APOE* genotypes ε2, ε3, and ε4 were determined by TaqMan genotyping of single nucleotide polymorphism (SNP) rs7412 and KASP genotyping of SNP rs429358. Genotyping was unsuccessful for one individual. Participants were categorized into those who carried at least one ε4 allele (*APOE* ε4) and those who did not (*APOE* ε4-).

Participants also self-reported their family history of dementia, i.e., whether a first-grade relative was affected by Alzheimer’s disease, vascular dementia or any other type of dementia. Two participants could not provide information about their family history (FH). The remaining participants were categorized into those with a positive FH (FH+) and those without (FH-).

### 2.3 Cognitive assessment

Immediate and 30 minutes delayed verbal and visual recall were assessed with the Rey Auditory Verbal Learning Test (RAVLT) (Rey, 1941; Schmidt, 1996) and the complex Rey figure Test (Rey, 1941). Short term topographical memory was measured with the Four Mountains Test (Chan et al., 2016). Spatial navigation was assessed with a virtual Morris Water Maze Task (Hamilton et al., 2002) where participants had to find and navigate to a hidden platform in a water pool. This task also included a motor control condition without a visible platform. Working memory capacity and executive functions were assessed with computerized tests from the Cambridge Brain Sciences battery (Hampshire et al., 2012; Owen et al., 2010). Working memory capacity was tested with digit and spatial span, distractor suppression with an adapted version of the Stroop test (Double-Trouble), problem solving with a version of the Tower of London task (the Tree task), abstract reasoning with grammatical reasoning and the odd-one-out task, as well as the ability to manipulate and organize spatial information with a self-ordered spatial span task. In addition, participants performed a paired-associate learning (PAL) and a choice-reaction time (CRT) task. Cognitive outcome measures were the number and latencies of correct responses as well as spatial navigation path length.

### 2.4 MRI data acquisition

MRI data were acquired on a 3T MAGNETOM Prisma clinical scanner (Siemens Healthcare, Erlangen, Germany). For hippocampal subfield segmentation and volumetric analyses, T_1_- and T_2_-weighted anatomical images were collected. T_1_-weighted images were acquired with a three-dimension (3D) magnetization-prepared rapid gradient-echo (MP- RAGE) sequence (256 x 256 acquisition matrix, TR = 2300 ms, TE = 3.06 ms, TI = 850ms, flip angle θ = 9°, 176 slices, 1mm slice thickness, FOV = 256 mm and acquisition time of ∼ 6 min). High resolution (0.4 x 0.4 x 2.5 mm voxel) T_2_-weighted anatomical images of the hippocampus were acquired with a turbo-spin-echo sequence in the coronal plane with TR = 3300 msec, TE = 84 msec, TI =, flip angle = 155°, 30 slices, 2.5mm slice thickness, FOV = 256 mm and acquisition time of ∼ 8 min.

High Angular Resolution Diffusion Imaging (HARDI) (Tuch et al., 2002) data (2 x 2 x 2 mm voxel) for the NODDI analyses were collected with a spin-echo echo-planar dual shell HARDI sequence with diffusion encoded along 90 isotropically distributed orientations (Jones et al., 1999) (30 directions at b-value = 1200 s/mm^2^ and 60 directions at b-value = 2400 s/mm^2^) and six non-diffusion weighted scans with dynamic field correction and the following parameters: TR = 9400ms, TE = 67ms, 80 slices, 2 mm slice thickness, FOV = 256 x 256 x 160 mm, GRAPPA acceleration factor = 2 and acquisition time of ∼15 min.

Quantitative magnetization transfer weighted imaging (qMT) data were acquired with an optimized 3D MT-weighted gradient-recalled-echo sequence (Cercignani and Alexander, 2006) to obtain magnetization transfer-weighted data with the following parameters: TR = 32 ms, TE = 2.46 ms; Gaussian MT pulses, duration t = 12.8 ms; FA = 5°; FOV = 24 cm, 2.5 x 2.5 x 2.5 mm^3^ resolution. The following off-resonance irradiation frequencies (Θ) and their corresponding saturation pulse nominal flip angles (ΔSAT) for the 11 MT-weighted images were optimized using Cramer-Rao lower bound optimization: Θ = [1000 Hz, 1000 Hz, 2750 Hz, 2768 Hz, 2790 Hz, 2890 Hz, 1000 Hz, 1000 Hz, 12060 Hz, 47180 Hz, 56360 Hz] and their corresponding ΔSAT values = [332°, 333°, 628°, 628°, 628°, 628°, 628°, 628°, 628°, 628°, 332°]. The longitudinal relaxation time, T_1_, of the system was estimated by acquiring three 3D gradient recalled echo sequence (GRE) volumes with three different flip angles (θ = 3°,7°,15°) using the same acquisition parameters as used in the MT-weighted sequence (TR = 32 ms, TE = 2.46 ms, FOV = 24 cm, 2.5 x 2.5 x 2.5 mm^3^ resolution). Data for computing the static magnetic field (B_0_) were collected using two 3D GRE volumes with different echo-times (TE = 4.92 ms and 7.38 ms respectively; TR= 330ms; FOV= 240 mm; slice thickness 2.5 mm) (Jezzard and Balaban, 1995).

### 2.5 Hippocampal subfield segmentations

Whole brain T_1_ and high resolution T_2_-weighted images were used as input images to segment subregions of the hippocampal formation with the FreeSurfer (version 6.0) (Iglesias et al., 2015) (Figure 1) and ASHS (Yushkevich et al., 2015b) hippocampal subfields segmentation tools. Detailed descriptions of the two pipelines are available on https://surfer.nmr.mgh.harvard.edu/fswiki/HippocampalSubfields and https://www.nitrc.org/projects/ashs.

**Figure 1.**
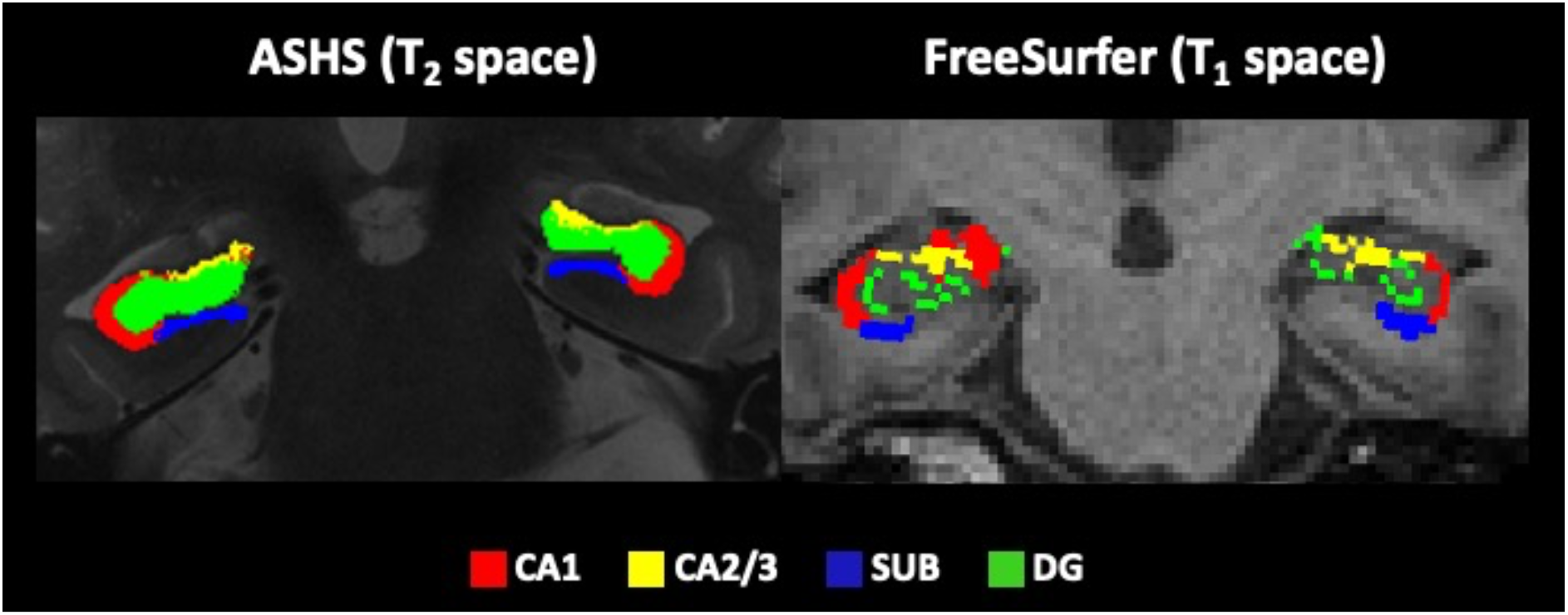
displays the hippocampal subfield regions that were automatically segmented from T_1_ and T_2_-weighted images using the Automated Segmentation of Hippocampal Subfields (ASHS) (Yushkevich et al., 2015b) and the FreeSurfer Segmentation of Hippocampal Subfields functionality (version 6) (Iglesias et al., 2015). Segmentations are shown for one participant with coronal images along the anterior-posterior hippocampal axis on a T_2_-weighted image for ASHS and a T_1_-weighted image for FreeSurfer. Abbreviations: cornu Ammonis 1 (CA1), CA2/3, subiculum (SUB) and dentate gyrus (DG).

In brief, the FreeSurfer version 6.0 cross-sectional segmentation pipeline firstly requires the fully automated analysis of the T1-weighted images with reconall (http://surfer.nmr.mgh.harvard.edu/) that involves skull stripping, correction for motion artefacts and field inhomogeneities, the registration of native data to and from the standard Talairach space, parcellation of cortical regions and the segmentation of subcortical structures including the hippocampi (Fischl, 2012). These processing steps were followed by a hippocampal subfield segmentation pipeline that utilises both T_1_ and T_2_-weighted images to identify anatomical landmarks for the bilateral segmentation of 12 subfields, i.e., CA1, CA2/3, CA4, subiculum, presubiculum, parasubiculum, molecular layer of the subiculum and CA fields, granule cell layer of the dentate gyrus, fimbria, hippocampus-amygdala transition area, hippocampal tail, and fissure (Iglesias et al., 2015). This pipeline is based on a statistical atlas of the hippocampus constructed from manual segmentation labels from both *in vivo* and high-resolution *ex vivo* data that were used to develop a Bayesian inference algorithm for the automatic segmentation of the hippocampus proper from T_1_ and T_2_-weighted structural images (Iglesias et al., 2015).

The ASHS software (https://sites.google.com/site/hipposubfields) implements a multi-atlas segmentation technique that involves registering the target MRI with a bank of T_2_-weighted atlas MRIs including manually labeled subregions. Here the ASHS UPenn PMC Atlas package was chosen for this purpose. Multi-atlas label fusion is then applied to select a consensus segmentation based on the shared similarity between target and atlas images. Systematic segmentation errors are minimized with a learning-based bias correction technique. Joint label fusion and corrective learning are then repeated by bootstrapping seeded by the segmentation results from the previous phase. ASHS leads to the segmentation of 10 region of interests, i.e., CA1, CA2, CA3, DG, subiculum, entorhinal cortex, Brodmann area 35, Brodmann area 36, collateral sulcus and miscellaneous regions. To allow meaningful extraction of lower resolution microstructural data, ASHS CA2 and CA3 regions were combined using the fslmaths utility from the Oxford Centre for Functional MRI of the Brain (FMRIB) Software Library (version 6.0) (Jenkinson et al., 2012).

For the purpose of comparing between the two segmentation techniques and for allowing meaningful extraction of lower resolution microstructural values, the present analysis focused on CA1, CA2/3, dentate gyrus and subiculum regions that were shown to be affected by aging and AD (de Flores et al., 2015a).

Mean hippocampal subfield and intracranial volumes (ICV) were extracted for each brain. Subfield volumes were then adjusted for ICV to correct for inter-individual differences in head size using the formula (subfield volume x 1000)/ ICV (from FreeSurfer or ASHS respectively).

### 2.6 HARDI and qMT data processing

A detailed description of the microstructural data processing has been provided (Metzler-Baddeley et al., 2019a; Metzler-Baddeley et al., 2019b). In brief, the dual-shell HARDI data were split and b = 1200 and 2400 s/mm^2^ data were corrected separately for distortions induced by the diffusion-weighted gradients and motion artifacts with appropriate reorientation of the encoding vectors (Leemans and Jones, 2009) in ExploreDTI (Version 4.8.3) (Leemans A et al., 2009). EPI-induced geometrical distortions were corrected by warping the diffusion-weighted image volumes to the T_1_ –weighted anatomical images (Irfanoglu et al., 2012). After preprocessing, the NODDI model (Zhang et al., 2012) was fitted to the HARDI data with the fast, linear model fitting algorithms of the Accelerated Microstructure Imaging via Convex Optimization (AMICO) framework (Daducci et al., 2015) to gain ISOSF, ICSF, and ODI maps.

Using Elastix (Klein et al., 2010), MT-weighted GRE volumes were co-registered to the MT-volume with the most contrast using a rigid body (6 degrees of freedom) registration to correct for inter-scan motion. Data from the 11 MT-weighted GRE images and T_1_-maps were fitted by a two-pool model using the Ramani pulsed-MT approximation (Ramani et al., 2002). This approximation provided MPF and *k_f_* maps. MPF maps were thresholded to an upper intensity limit of 0.3 and *k_f_* maps to an upper limit of 3.0 using the FMRIB’s fslmaths imaging calculator to remove voxels with noise-only data.

All microstructural maps were spatially aligned with the hippocampal subfield masks by co-registration with the T_1_-weighted anatomical space as reference image with linear affine registration (12 degrees of freedom) using FMRIB’s Linear Image Registration Tool (FLIRT). Spatial alignment of microstructural maps to ASHS hippocampal subfield masks involved an additional warping to the T_2_-weighted space with FLIRT.

### 2.7 Statistical analysis

Statistical analyses were conducted in SPSS version 26 (IBM, 2011). All data were examined for normal distribution and for outliers, defined as above or below three times the interquartile range (75th percentile value - 25th percentile value).

*Missing data*: FreeSurfer hippocampal subfield segmentations could be performed for all 158 datasets and ASHS segmentations for a total of 153 datasets. For FreeSurfer one volume and for ASHS two volume measurements were excluded as outliers. For the microstructural data, 1% of the data were excluded as outliers for FreeSurfer and 5% for ASHS segmentations.

For the principal component analysis (PCA), each participant’s volumetric and microstructural data for the 16 hippocampal subfield segmentations [2 (FreeSurfer, ASHS) x 2 (left, right) x 4 (CA1, CA2/3, DG, subiculum)] were concatenated to form n = 2528 observations. The dependent variables were represented by the seven brain measurements (ICV-adjusted volumes, MPF, *k_f_*, R_1_, ODI, ISOSF, ICSF). Bartlett’s test of sphericity and the Kaiser-Meyer-Olkin (KMO) test were used to check that the data were suitable for PCA [KMO = 0.57, Chi^2^ (21) = 2842.2, p < 0.001]. PCA was then carried out using a procedure with orthogonal Varimax rotation of the component matrix. Components were extracted based on the Kaiser criterion of including all components with an eigenvalue > 1 (IBM, 2011), by inspecting Cattell’s scree plots (Cattell, 1952) and by assessing each component with regards to their interpretability. Component loadings that exceeded a value of 0.5 were considered as significant. The dimensionality of the cognitive data for all 158 participants was also reduced with PCA using the same procedure as described above [KMO = 0.61, Chi^2^ (630) = 2463.37, p < 0.001].

Each participant’s PCA least squares regression component scores (DiStefano et al., 2009) were subsequently entered as dependent variables in a multivariate analysis of covariance (MANCOVA) that tested for main and interaction effects of the risk factors *APOE* genotype (ε4+, ε4-), FH (FH+, FH-), and WHR (WHR+, WHR-) as well as for the effects of the segmentation protocol (FreeSurfer, ASHS) and hippocampal subfield segmentations (bilateral CA1, CA2/3, DG, subiculum). Age, sex and IQ-scores from the NART-R (Nelson, 1991) were included as covariates. Similarly, MANCOVA tests for risk effects on cognitive component scores, while controlling for age, sex, and IQ, were completed.

Significant omnibus effects were further investigated with post-hoc comparisons using univariate analysis of covariance (ANCOVA) and independent t-tests. Relationships between brain structure and cognitive component scores were studied using hierarchical linear regression models that first entered as independent variables age and sex, and then in a second model volumetric and microstructural measurements of CA1, CA2/3, DG and subiculum in a stepwise fashion to predict the variance in the cognitive component data. To reduce the number of independent variables entered into the regression model and hence model overfit, macro- and microstructural measurements were averaged over protocol and hemisphere for each subfield.

First and post-hoc models were corrected for multiple comparisons with a False Discovery Rate (FDR) of 5% using the Benjamini-Hochberg procedure (Benjamini and Hochberg, 1995). The 5% FDR was applied to all statistical tests that related to the same theoretical inference (Lakens, 2014).

All reported p-values, unless stated otherwise, were Benjamini-Hochberg adjusted (p_BHadj_) and two-tailed. Information about effects sizes was provided with the partial eta squared index η_p_^2^ for MANCOVA analyses and R^2^ for regression analyses.

## 3. Results

### 3.1 Principal Component Analysis (PCA) of brain measurements

PCA extracted three components that explained 67% of the variance in the hippocampal macro- and microstructural data (Table 2). The first component explained 28.2% of the data variance and had positive loadings > 0.5 from MPF, R_1_ and ICSF sensitive to myelin and neurite packing. The second component explained an additional 24.5% of the variance and had positive loadings > 0.5 from ODI and ICV-adjusted volumes and a negative loading from ISOSF and thus captured tissue size and complexity. The third component explained 14.3% of variance and had a high loading from *k_f_*, that may reflect tissue metabolism.

**Table 2.**
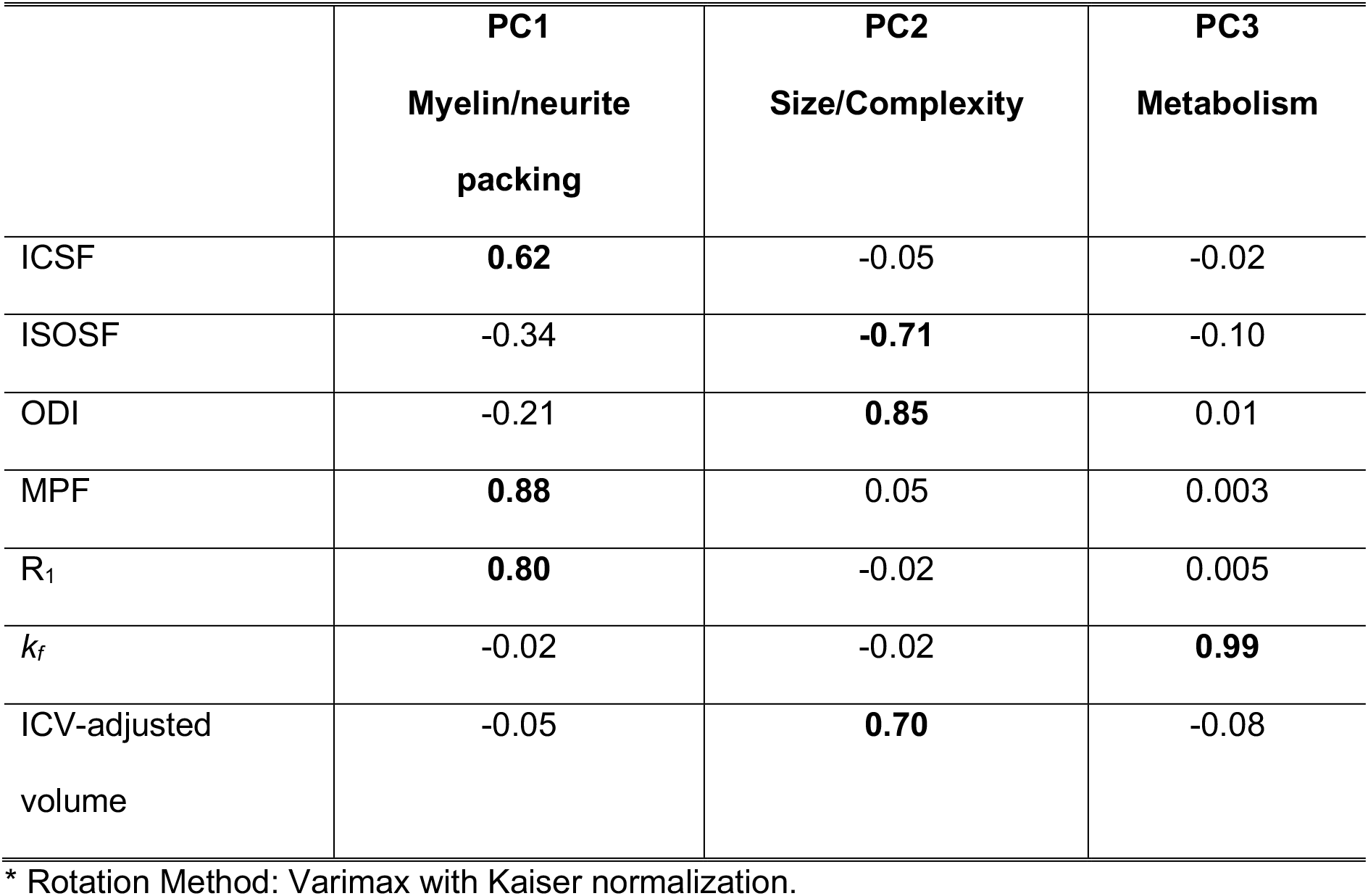
Rotated Component Matrix of the Principal Component Analysis within the macro- and microstructural data from FreeSurfer and ASHS hippocampal subfields (N = 2528).

### 3.2 Multivariate Analysis of Covariance (MANCOVA) of macro- and microstructural PCs

*Omnibus effects*: There were significant main effects for age [F(3,2124) = 13.72, p_BHadj_ < 0.001, η_p_^2^ = 0.02], sex [F(3,2124) = 22.24, p_BHadj_ < 0.001, η_p_^2^ = 0.03], WHR [F(3,2124) = 6.8, p_BHadj_ < 0.001, η_p_^2^ = 0.01], protocol [F(3,2124) = 128.7, p_BHadj_ < 0.001, η_p_^2^ = 0.15] and hippocampal subfield [F(9,6378) = 322.2, p_BHadj_ < 0.001, η_p_^2^ = 0.3]. Significant interaction effects were present between protocol and hippocampal subfield [F(9,6378) = 111.29, p_BHadj_ < 0.001, η_p_^2^ = 0.14] and between *APOE* and WHR [F(3,2124) = 5.51, p_BHadj_ = 0.005, η_p_^2^ = 0.008].

*Post hoc effects*: Protocol and hippocampal subfield had significant effects on the myelin/neurite packing PC1 [Protocol: F(1,2126) = 21.5, p_BHadj_ < 0.001, η_p_^2^ = 0.01; Subfield: F(3,2126) = 236.7, p_BHadj_ < 0.001, η_p_^2^ = 0.25] and the size/complexity PC2 [Protocol: F(1,2126) = 381.7, p_BHadj_ < 0.001, η_p_^2^ = 0.15; Subfield: F(3,2126) = 1483.2, p_BHadj_ < 0.001, η_p_^2^ = 0.68]. Overall ASHS relative to FreeSurfer segmentations had smaller PC1 values [t(2248) = -3.9, p_BHadj_ < 0.001] (Figure 2 A) but larger PC2 values [t(2248) = 11.2, p_BHadj_ < 0.001] (Figure 2B). With regards to the subfields, subiculum was associated with the largest PC1 myelin/neurite packing values (Figure 2C), while CA2/3 had the lowest PC2 size/complexity values (Figure 2D). No effects were observed for the PC3 Metabolite component.

**Figure 2.**
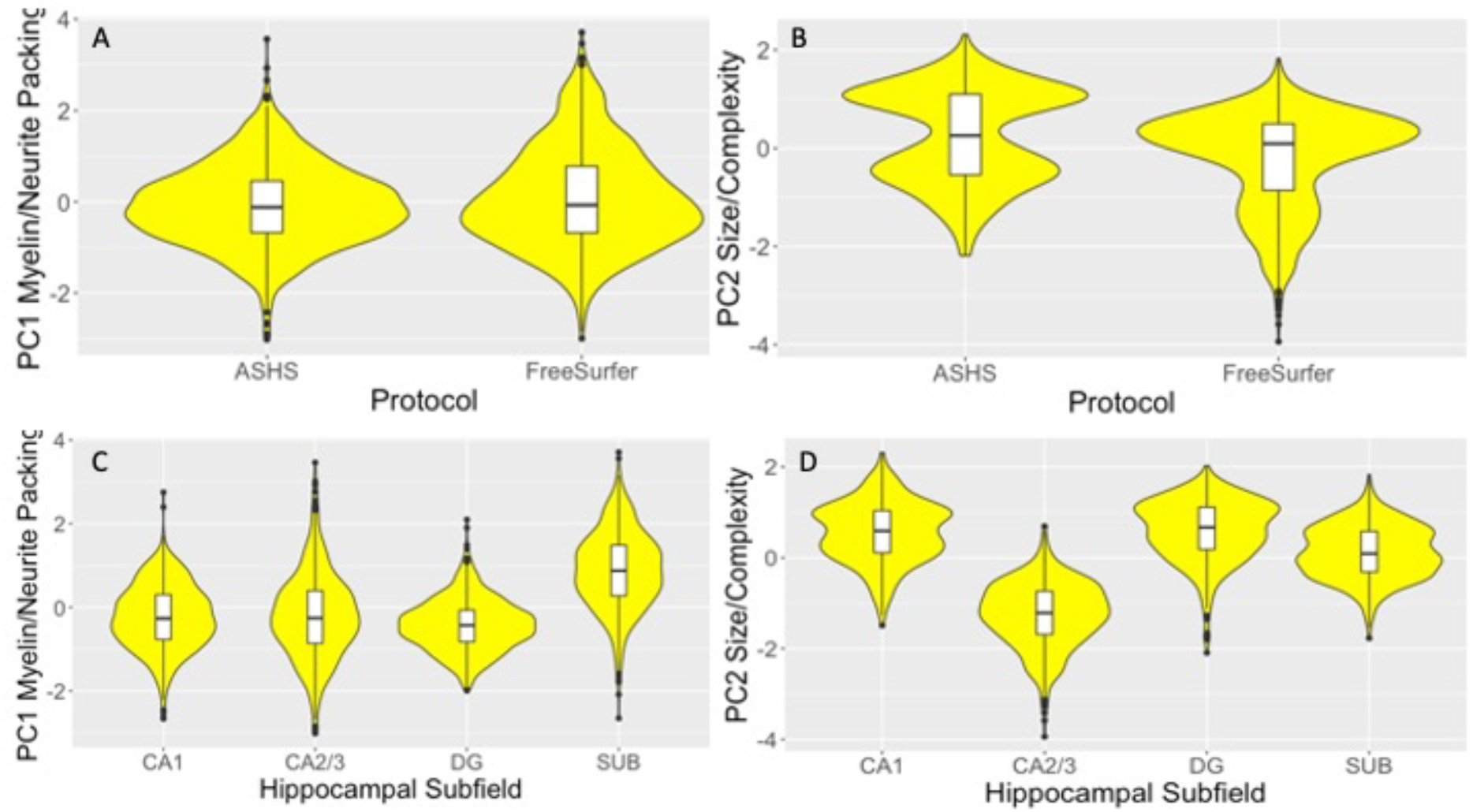
Violin plots with overlaid box plots of the difference between; A) the Automatic Segmentation of Hippocampal Subfields (ASHS) and the FreeSurfer (version 6) segmentation protocols in the myelin/neurite packing principal component (PC); B) in the size/complexity PC, C) between the hippocampal subfields cornu Ammonis (CA) 1, CA2/3, dentate gyrus (DG) and subiculum in the myelin/neurite packing principal component (PC); D) in the size/complexity PC. Boxplots display the median and the interquartile range and violin plots the kernel probability density, i.e., the width of the yellow area represents the proportion of the data located there.

Furthermore, protocol and subfields interacted with each other in both components [PC1: F(3,2126) = 37.1, p_BHadj_ < 0.001, η_p_^2^ = 0.05; PC2: F(3,2126) = 385.7, p_BHadj_ < 0.001, η_p_^2^ = 0.35] (Figure 3). FreeSurfer compared with ASHS segmentations were associated with larger PC1 myelin/neurite packing values, primarily due to larger values in the subiculum (Figure 3A). In contrast, ASHS compared with FreeSurfer segmentations showed larger PC2 size/complexity values (Figure 3B), due to larger estimations in CA1, CA2/3 and DG but not subiculum.

**Figure 3.**
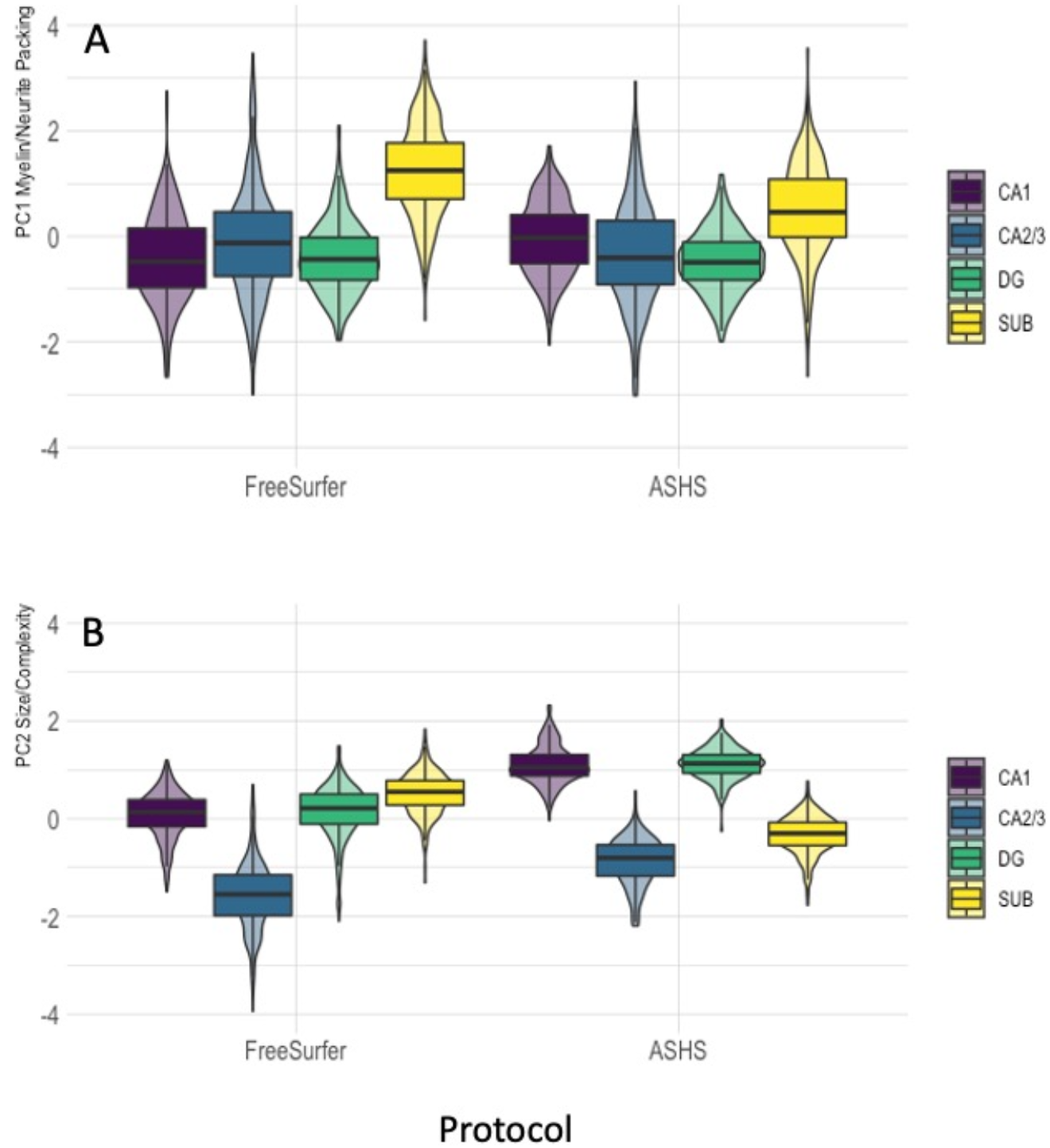
Violin plots with overlaid box plots displaying the effects of protocol as a function of hippocampal subfields on; A) the myelin/neurite packing principal component; B) the size/complexity component. Boxplots display the median and the interquartile range and violin plots the kernel probability density. ASHS = Automated Segmentation of Hippocampal Subfields, CA = cornu Ammonis, DG = dentate gyrus, SUB = subiculum.

Age had a significant effect on the myelin/neurite packing PC1 [F(1,2126) = 37.3, p_BHadj_ < 0.001, η_p_^2^ = 0.02]. More specifically, age was associated with a reversed U-shape in PC1 values having a peak in the forties, with youngest and oldest participants showing the lowest values (Figure 4A). Sex had an effect on the size/complexity PC2 [F(1,2126) = 56.3, p_BHadj_ < 0.001, η_p_^2^ = 0.03], with males showing reduced PC2 values compared with females [t(2279) = 5.3, p_BHadj_ < 0.001] (Figure 4B).

**Figure 4.**
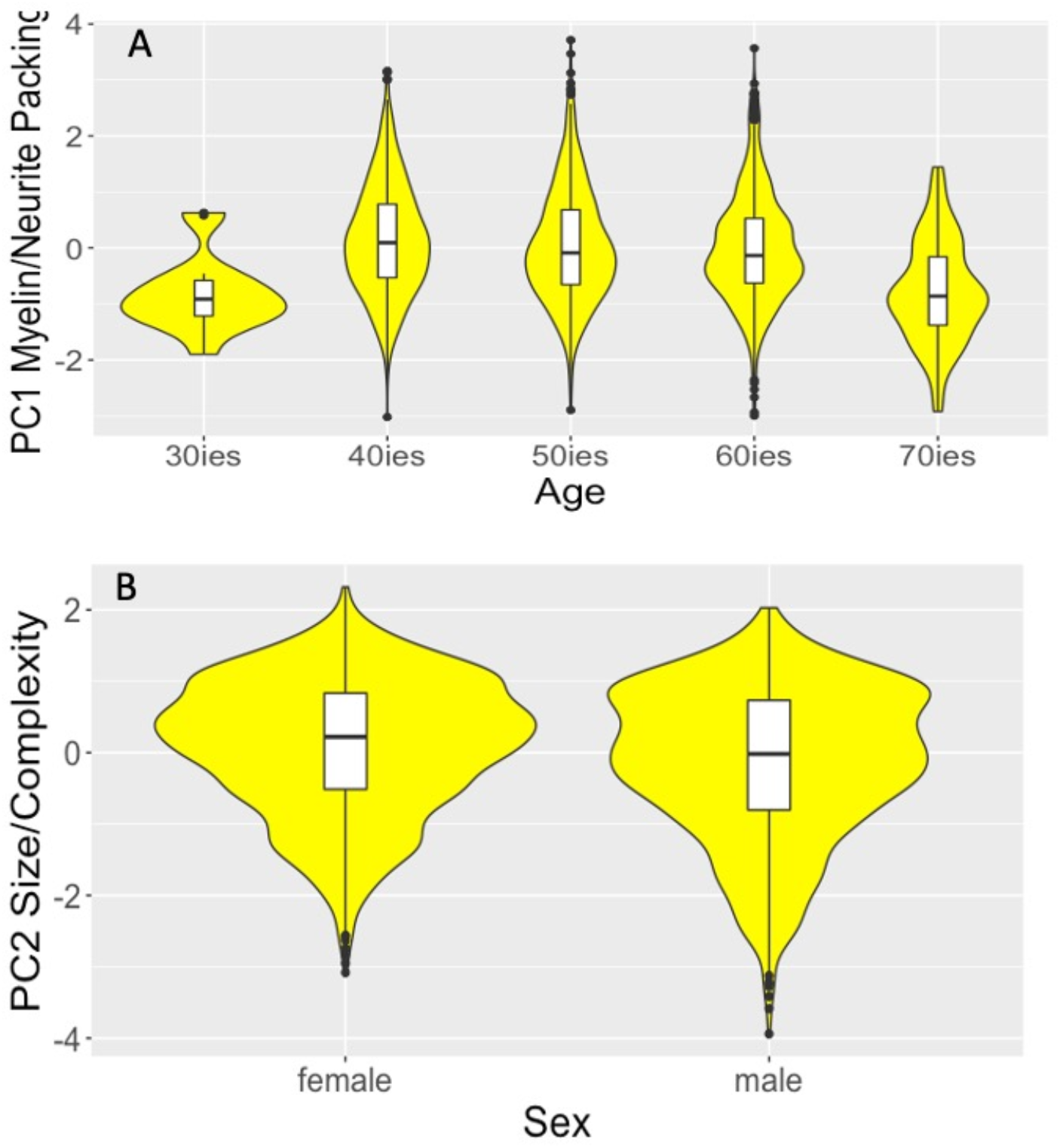
Violin plots with overlaid box plots displaying the effects of A) age on the myelin/neurite packing principal component (PC) and B) of sex on the size/complexity PC. Boxplots display the median and the interquartile range and violin plots the kernel probability density.

WHR affected myelin/neurite packing and size/complexity components [PC1: F(1,2126) = 11.6, p_BHadj_ = 0.002, η_p_^2^ = 0.005; PC2: F(1,2126) = 5.5, p_BHadj_ = 0.036, η_p_^2^ = 0.003] as centrally obese individuals relative to those with a normal WHR had lower values in both components [PC1: t(2231) = 4.14, p_BHadj_ < 0.001; PC2: t(2231) = 4.15, p_BHadj_ < 0.001] (Figure 5). Moreover, WHR interacted with *APOE* on size/complexity [F(1,2126) = 15.2, p_BHadj_ < 0.001, η_p_^2^ = 0.007] (Figure 6). Whilst *APOE* ε4*-*individuals exhibited obesity-related size/complexity reductions [t(1341) = 4.14, p_BHadj_ < 0.001], this was not the case for *APOE* ε4*+* individuals [t(880) = 1.6, p= 0.1] (Figure 6).

**Figure 5.**
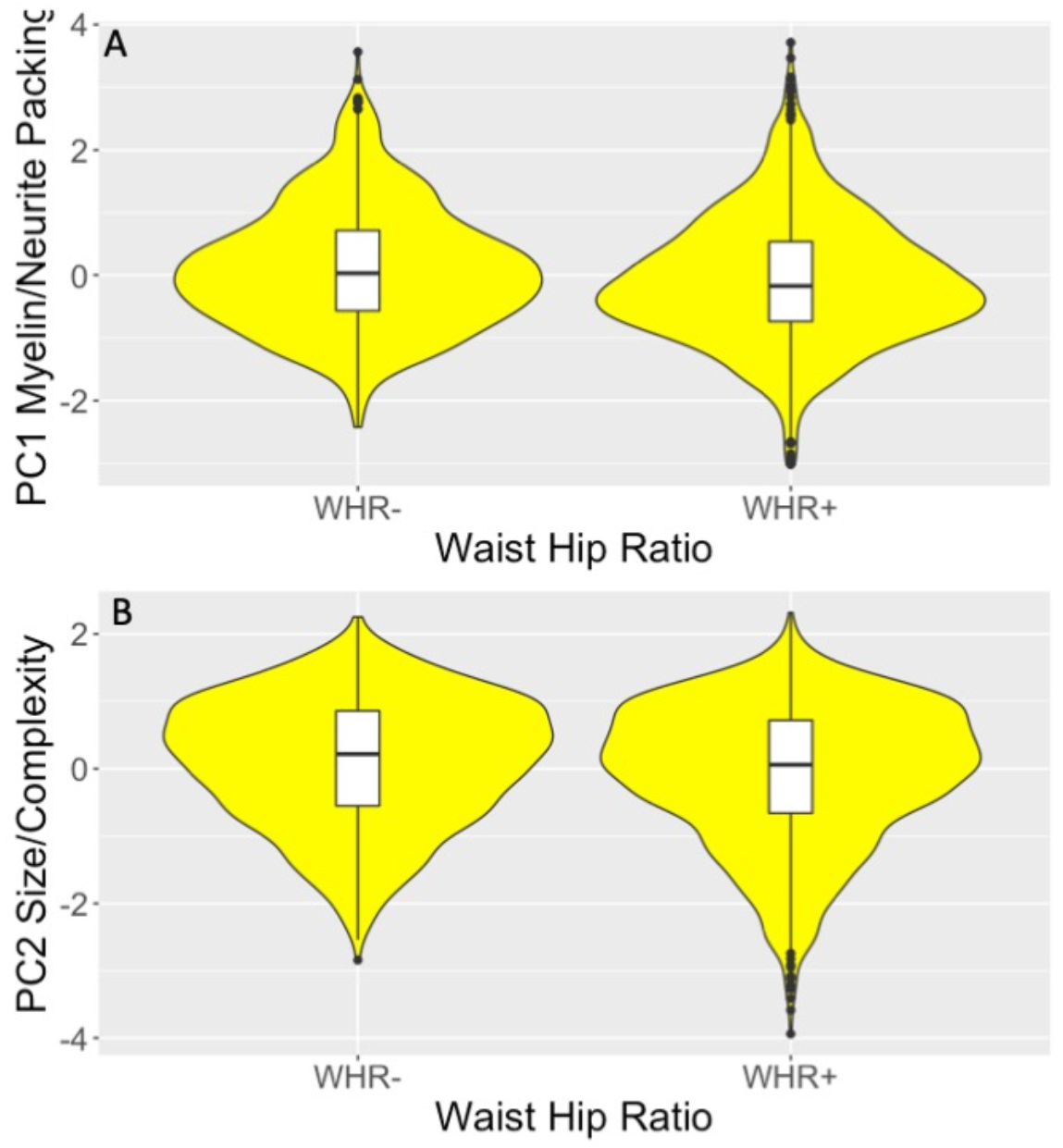
Violin plots with overlaid box plots displaying the effects of Waist-Hip-Ratio (WHR); A) the myelin/neurite packing principal component (PC); B) the size/complexity PC. Boxplots display the median and the interquartile range and violin plots the kernel probability density.

**Figure 6.**
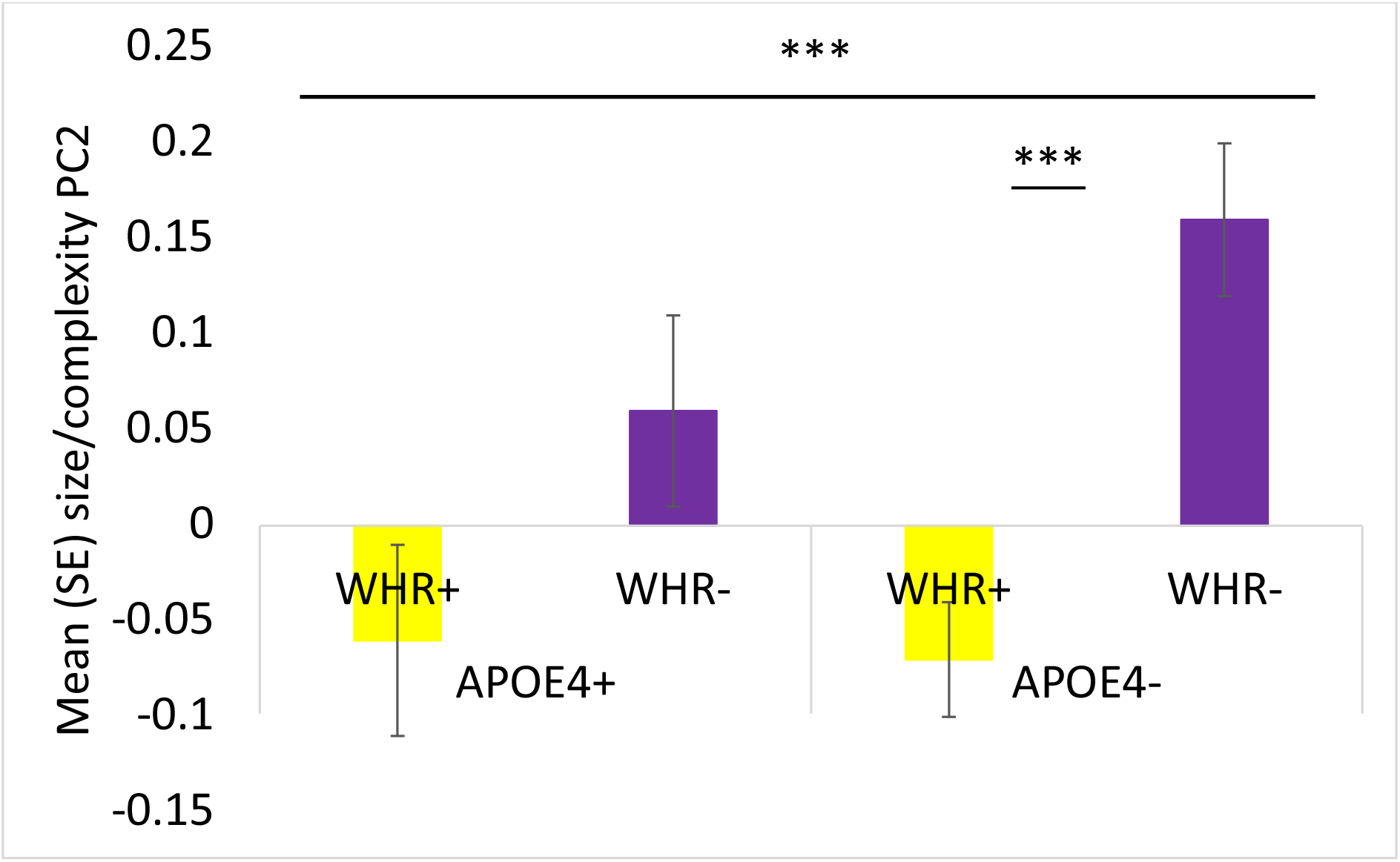
Column figure displaying the mean and standard errors of the size/complexity principal component (PC) as a function of Apolipoprotein E (*APOE*) genotype and Waist-Hip-Ratio (WHR). APOE4+ = *APOE* ε4-carriers, APOE4- = *APOE* ε4-non-carriers; WHR+ = individuals with WHR in abdominal overweight/obese range; WHR- = individuals with WHR in healthy range. *** p_BHadj_ < 0.001

### 3.3 Exploring the pattern of age and risk effects across hippocampal subfields

To explore the patterns of age and risk effects across the four hippocampal subfields CA1, CA2/3, DG, and subiculum, separate ANCOVAs were carried out on the PC data for each subfield concatenated across hemisphere and protocol.

Significant age effects (controlled for sex and NART-R IQ) on the myelin/neurite packing PC1 were observed for CA1 [F(4,564) = 12.9, p_BHadj_ < 0.001, η_p_^2^ = 0.08], for CA2/3 [F(4,567) = 5.7, p_BHadj_ < 0.001, η_p_^2^ = 0.04], for DG [F(4,556) = 7.4, p_BHadj_ < 0.001, η_p_^2^ = 0.05] and subiculum [F(4,568) = 11.7, p_BHadj_ < 0.001, η_p_^2^ = 0.08].

Trends for WHR effects (controlled for age, sex and NART-R IQ) on the size/complexity PC2 were present in CA1 (p_BHadj_ = 0.06), CA2/3 (p_BHadj_ = 0.06) and DG (p_BHadj_ = 0.09) but not for the subiculum (p = 0.6). Trends for interaction effects between *APOE* and WHR were present for CA1 (p = 0.06) and DG (p = 0.06) but not for CA2/3 (p = 0.32) or subiculum (p = 0.6).

### 3.4 PCA of the cognitive data

Four principal components were extracted that explained together 44% of the variance in the cognitive data (Table 3). The first PC explained 15.2% of the variance and had high loadings > 0.5 from all RAVLT measures and hence reflected verbal recall performance. The second PC explained 13.5% of the variance with loadings from first move latencies in the virtual spatial navigation task. The third PC explained an additional 9.5% of the variance with loadings from spatial navigation path length. The fourth PC explained 5.8% of data variance and had loadings from immediate and delayed recall of the Rey figure, hence reflecting visual recall abilities.

**Table 3.** Rotated component matrix of the principal component analysis of the cognitive performance

| Cognitive Tests | Components |  |  |  |
| --- | --- | --- | --- | --- |
|  | Verbal<br>Memory | Spatial<br>Navigation<br>First Move | Spatial<br>Navigation<br>Path Length | Visual<br>Memory |
| RAVLT List A 1<sup>st</sup> IR | **0.78** | 0.01 | -0.02 | 0.06 |
| RAVLT List A 2nd IR | **0.85** | 0.02 | -0.03 | 0.01 |
| RAVLT List A 3rd IR | **0.84** | 0.09 | -0.01 | -0.03 |
| RAVLT List A 4th IR | **0.81** | 0.02 | -0.05 | -0.17 |
| RAVLT List A 5th IR | **0.75** | 0.03 | -0.01 | -0.08 |
| RAVLT List B 1<sup>st</sup> IR | **0.47** | -0.25 | -0.06 | 0.06 |
| RAVLT List A 6th RaD | **0.81** | -0.02 | 0.07 | 0.13 |
| RAVLT List A DR | **0.81** | 0.01 | 0.02 | 0.14 |
| Four Mountains Test | 0.23 | -0.10 | -0.23 | **0.50** |
| Rey Copy | 0.13 | -0.13 | 0.06 | 0.43 |
| Rey Figure IR | -0.02 | -0.09 | -0.02 | **0.80** |
| Rey Figure DR | 0.02 | -0.03 | -0.04 | **0.85** |
| Digit Span | 0.13 | -0.27 | -0.09 | 0.001 |
| Spatial Span | -0.14 | -0.06 | -0.23 | 0.30 |
| Double Trouble | -0.01 | -0.16 | -0.21 | 0.19 |
| Tree Task | 0.07 | -0.04 | -0.09 | -0.21 |
| Odd one out | -0.07 | -0.16 | -0.02 | -0.03 |
| Paired Associate Learning | 0.09 | -0.11 | -0.25 | -0.20 |
| Self-ordered search | -0.28 | 0.02 | -0.11 | -0.07 |
| Grammatical Reasoning | 0.01 | -0.29 | -0.02 | 0.06 |
| Choice Reaction Time | -0.01 | 0.41 | 0.15 | -0.34 |
| Spatial Navigation: |  |  |  |  |
| FM Block 2 | 0.35 | **0.75** | -0.07 | -0.03 |
| FM Block 3 | 0.01 | **0.78** | -0.15 | 0.03 |
| FM Block 4 | -0.05 | **0.76** | -0.22 | -0.02 |
| FM Block 5 | 0.01 | **0.67** | -0.40 | 0.01 |
| FM Block 6 | -0.2 | **0.74** | 0.03 | -0.03 |
| TL Block 2 | -0.04 | **0.63** | 0.44 | -0.06 |
| TL Block 3 | 0.18 | 0.35 | 0.32 | -0.22 |
| TL Block 4 | 0.12 | 0.41 | 0.45 | -0.25 |
| TL Block 5 | -0.23 | 0.43 | **0.57** | -0.16 |
| TL Block 6 | -0.11 | **0.66** | 0.09 | -0.30 |
| PL Block 2 | -0.02 | -0.001 | **0.68** | 0.08 |
| PL Block 3 | 0.22 | -0.22 | **0.58** | -0.02 |
| PL Block 4 | 0.15 | -0.14 | **0.71** | -0.09 |
| PL Block 5 | -0.16 | -0.08 | **0.80** | -0.03 |
| PL Block 6 | -0.16 | -0.06 | 0.09 | -0.45 |

Loadings > 0.5 are highlighted in bold. Abbreviations: DR = delayed recall, IR = immediate recall, RaD = recall after distraction, RAVLT = Rey Auditory Verbal Learning Test.
<sup>a</sup> Rotation method: Varimax with Kaiser normalization.

### 3.5 MANCOVA of cognitive PCs

There were significant omnibus effects of age [F(4,99) = 7.2, p_BHadj_ < 0.001, η_p_^2^ = 0.23] and sex [F(4,99) = 22.24, p_BHadj_ =0.02, η_p_^2^ = 0.14]. Age affected first move latencies in the spatial navigation task [F(1,102) = 19.6, p_BHadj_ < 0.001, η_p_^2^ = 0.16] such that latencies increased with age (r = 0.38, p < 0.001). Sex had an effect on verbal recall performance [F(1,102) = 10.6, p_BHadj_ = 0.008, η_p_^2^ = 0.16] with women performing better in the RAVLT than men. There were no effects of risk.

### 3.6 Regression analysis of brain-cognition relationships

Table 4 summarizes the results of the linear hierarchical regression analyses. As age and sex had significant effects on the cognitive components, they were first entered as independent variables prior to testing for the effects of hippocampal subfield macro- and microstructure. For the PC4 Visual Recall, 16% of the data were explained by a final model that included contributions from the CA fields, i.e., CA2/3 ICSF, CA1 volume and CA1 ISOSF. In addition, 17% of the variance in the PC2 Spatial Navigation First Move Latencies was explained by age and sex. The regression models for the PC1 and PC3 were not significant.

**Table 4.**
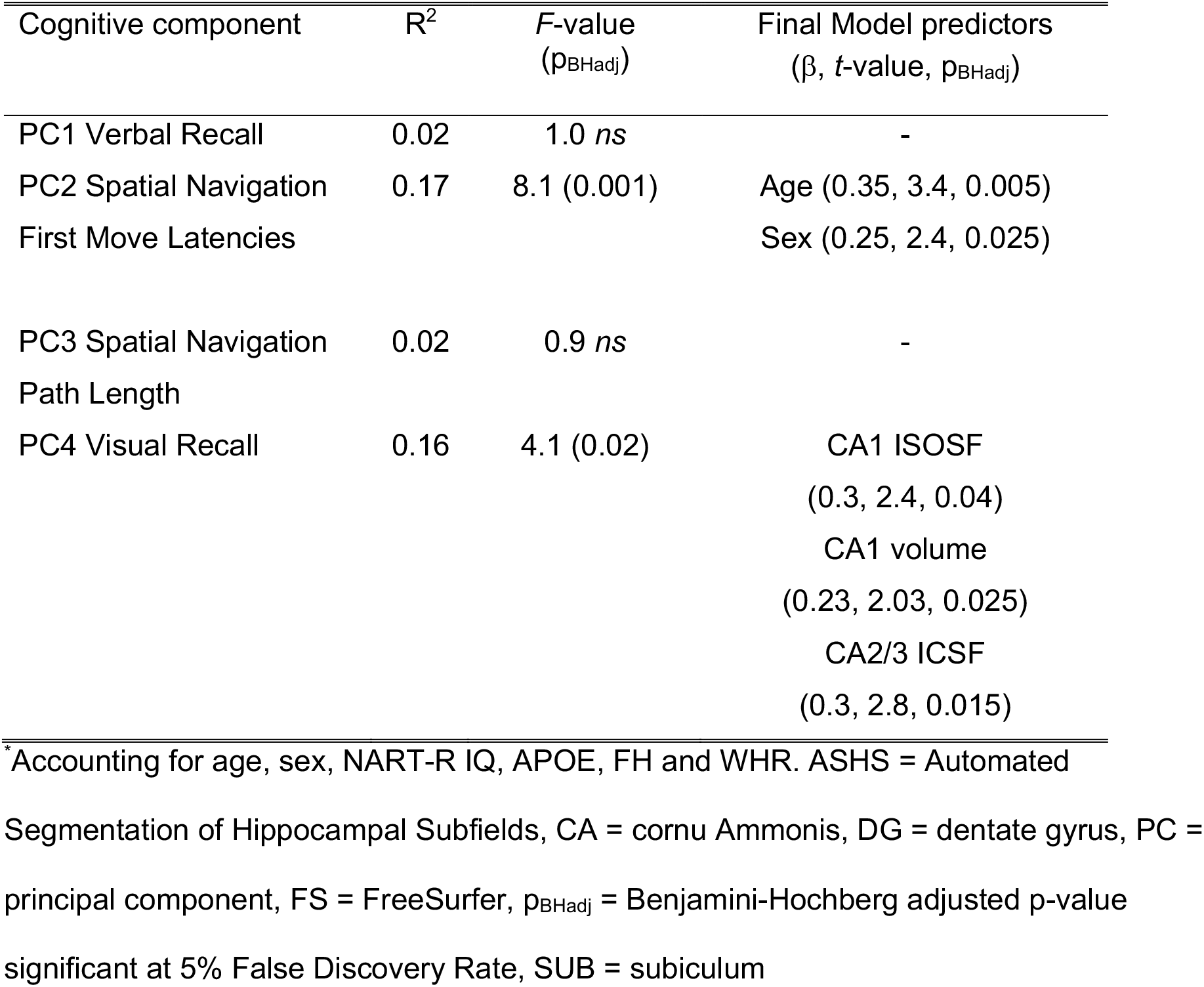
Results of hierarchical stepwise regression analyses testing first for the effects of age, sex, verbal intelligence and secondly for the effects of macro- and microstructural measurements from CA1, CA2/3, Dentate Gyrus (DG) and subiculum on cognitive components

## 4. Discussion

Dissociating the effects of healthy aging on hippocampal subfields from those related to genetic and lifestyle risk of dementia could be key to effectively targeting interventions for older-age memory impairments. Thus, the primary objective of the present study was to investigate age and age-independent effects of three major risk factors, i.e., carriage of the APOE ε4 genotype, a positive FH of dementia, and central obesity, on the macro- and microstructure of the hippocampal formation in a sample of 158 cognitively healthy adults.

We characterized properties of the hippocampal formation with subfield volumes based on T_1_ and high-resolution T_2_-weighted images as well as with microstructural measurements from NODDI and qMT imaging to gain complementary information about apparent myelin, neurite packing, free water, and metabolism. Accounting for overlapping information between the various MRI measurements, PCA was employed to reduce the data dimensionality to three principal components that reflected myelin/neurite packing, size/complexity, and metabolic gray matter properties. These components were then investigated across the main subfields of the hippocampal formation, i.e., CA1, CA2/3, DG, and subiculum, which were shown to be particularly vulnerable to the impact of aging and disease (de Flores et al., 2015b). Subfields were segmented using two widely employed and freely available, automated segmentation protocols, i.e., ASHS and FreeSurfer (version 6). This allowed us to assess any potential interaction effects between the type of protocol and risk factors on the analysis of hippocampal subfield properties.

While type of labelling protocol and hippocampal subfields were associated with absolute differences in component measures, they did not interact with the risk factors, suggesting that risk effects did not significantly differ between ASHS and FreeSurfer segmentations. Overall ASHS segmentations resulted in larger size/complexity but lower myelin/neurite packing estimates (Figure 2). ASHS segmentations of CA1, CA2/3 and DG were larger than those of FreeSurfer while subiculum labels were larger for FreeSurfer than ASHS (Figure 3). Myelin/neurite packing signals were largest in the subiculum for both protocols but particularly so for FreeSurfer (Figure 3). Both ASHS and FreeSurfer CA2/3 labels were the smallest subfields while CA1 and DG were comparable in size/complexity. The subiculum was smaller than CA1 and CA2/3 for ASHS but larger for FreeSurfer.

These component differences were caused by disagreements in the anatomical labels between the two segmentation protocols that are known to be most pronounced at the CA1/subiculum boundary and within the anterior portion of the hippocampal formation (Yushkevich et al., 2015a). In addition, both protocols differed in the region of the hippocampal formation that was labelled and the number of subfield segmentations. While FreeSurfer labels 12 regions (CA1, CA2/3, CA4, subiculum, presubiculum, parasubiculum, molecular layer of the subiculum and CA fields, granule cell layer of the dentate gyrus, fimbria, hippocampus-amygdala transition area, hippocampal tail, and fissure) (Iglesias et al., 2015), ASHS segments ten regions (CA1, CA2, CA3, DG, subiculum, entorhinal cortex, Brodmann area 35, Brodmann area 36, collateral sulcus and miscellaneous regions) (Yushkevich et al., 2015b). As many of these subfields were too small to extract meaningful lower resolution microstructural information, we focused on the analysis of the four main subfields of the hippocampal formation that were labelled by both protocols (CA1, CA2/3, DG and subiculum).

It should also be noted that the final ASHS segmentation outputs were in high resolution T_2_-weighted space while those for FreeSurfer were in T_1_-weighted space (Figure 1) suggesting differences in the use of multi-spectral information between the pipelines. A recent study found significant variations in FreeSurfer volume estimations of hippocampal subfields depending on the input images (T_1_ and/or T_2_) (Seiger et al., 2021). It is therefore likely that differences in the use of standard T_1_ and high-resolution T_2_-based information may have contributed to the discrepancies observed between the two protocols. Importantly though for our primary research question, despite these significant differences between the ASHS and FreeSurfer labels, there were no interaction effects between the type of protocol and any of the three risk factors, suggesting that a comparable risk pattern was observed across both protocols.

With regards to risk factors, we observed that central obesity as measured with the WHR was associated with reductions in the myelin/neurite packing and the size/complexity but not the metabolic component (Figure 5). These findings are consistent with accumulating evidence that obesity is associated with adverse effects on the hippocampus and memory functions (Anan et al., 2010; Dekkers et al., 2019; Khan, et al., 2015; Mueller et al., 2012; Spencer et al., 2017; Stranahan, 2015; Willette and Kapogiannis, 2015). For instance, a recent analysis of data from 12,000 participants (45-76 years of age) of the UK Biobank study, reported that total body fat was related to smaller subcortical gray matter volumes including the hippocampus in men (Dekkers et al., 2019). This study also reported obesity-related differences in whole brain white matter microstructure measured with fractional anisotropy and mean diffusivity. Similarly, previous analyses of CARDS data found WHR to be positively correlated with hippocampal atrophy [as estimated with the free water signal (ISOSF)] and negatively with fornix apparent myelin MPF and *k_f_* (Metzler-Baddeley et al., 2019a). In this study, WHR was a close estimate of abdominal visceral but not subcutaneous fat fractions while Body Mass Index (BMI) estimated subcutaneous but not visceral fat (Metzler-Baddeley et al., 2019a). As BMI had no effects on brain microstructure, these findings suggested that the correlations between WHR and hippocampal and fornix microstructure were driven by excessive visceral rather than body fat per se, consistent with accumulating evidence for visceral fat-related adverse effects on the hippocampus, brain white matter, and mortality (Anan et al., 2010; Koster et al., 2015; Koster and Schaap, 2015; Koster et al., 2010; Lampe et al., 2019). Comparable to Dekkers et al. (2019), men were more centrally obese and had higher visceral fat fractions and higher hippocampal ISOSF (Metzler-Baddeley et al., 2019a).

Central obesity, notably excessive visceral fat, are associated with multiple metabolic alterations affecting blood cholesterol, glucose, and insulin levels, that can lead to cardio- and cerebrovascular disease and Type 2 diabetes (Dommermuth and Ewing, 2018; Whitmer et al., 2007). Central obesity at midlife may also be accompanied with systemic, low-grade inflammation (Cox et al., 2015; Guillemot-Legris and Muccioli, 2017). Diet-induced obesity in animal studies has been shown to trigger inflammation in the hippocampus, which in turn impaired synaptic functioning and spatial memory (Hao et al., 2016). In humans, individuals with a genetic polymorphism associated with pro-inflammatory state and AD risk had smaller CA1-2, CA3-DG and subiculum than healthy controls (Raz et al., 2015). All of these obesity-related metabolic changes in lipid, glucose and immune responses are thought to contribute to the risk of developing dementia in older age (Businaro et al., 2018; Profenno et al., 2010; Ricci et al., 2017).

Furthermore, it is also increasingly recognized that obesity may interact with genetic risk factors, notably *APOE* ε4 (Jones and Rebeck, 2018; Mole et al., 2020; Zade et al., 2013). Indeed, here we observed interaction effects between *APOE* and WHR on the size/complexity of the hippocampal formation. While all obese individuals showed reduced size/complexity, this effect was only significant for APOE ε4 non-carriers (p < 0.001) but not for APOE ε4 carriers (p = 0.1) (Figure 6). As can be seen in Figure 6, APOE ε4 carriers did not show a significant obesity effect because size/complexity was attenuated in normal-weighted APOE ε4 carriers. Thus central obesity appeared to be related to atrophy in the hippocampal formation regardless of an individual’s *APOE* or FH status (as no effects of FH were present). However, *APOE* ε4 carriage alone appeared to be associated with adverse effects on the hippocampus that may have masked any metabolic and vascular benefits of a healthy WHR. Or expressed differently, keeping a healthy weight may not be sufficient to compensate for *APOE* ε4-driven hippocampal atrophy. These risk effects were present for data collapsed across all hippocampal subfields. There was no evidence for any subfield specific vulnerability to the impact of obesity and *APOE*. However, it should be noted that we observed trends for CA1, CA2/3 and DG but not the subiculum. Future larger studies are required to clarify whether these regions are particularly susceptible to obesity and *APOE* ε4 related tissue changes.

A previous analysis of data from the Framingham Offspring cohort also highlighted complex synergistic effects between *APOE* and obesity (Zade et al., 2013). They found several *APOE-*related modifications of correlations between individual differences in WHR on one hand and brain structure and cognition on the other. For instance, *APOE* ε4 carriers showed stronger negative relationships between WHR and executive and memory functions as well as larger correlations between WHR and white matter hyperintensities. Interestingly, they also reported a stronger correlation between WHR and frontal brain volume for *APOE* ε4 non-carriers similar to our observations here.

In addition, it is likely that obesity effects will be modulated by other polygenic risk factors than *APOE* ε4 (Woo and Reifman, 2018). While the CARDS sample was too small to quantify AD and/or obesity-related polygenic risk hazards (Escott-Price et al., 2014; Escott-Price et al., 2015), we included family history of dementia as a variable that captures environmental and genetic risk factors beyond *APOE* ε4. In the present analysis of hippocampal gray matter microstructure we did not find any effect of FH, but previously we observed widespread interaction effects between *APOE*, FH, and WHR on white matter microstructure, that were most pronounced in the right parahippocampal cingulum (Mole et al., 2020). More specifically, *APOE* ε4 carriers with a positive FH, showed obesity-related reductions in apparent myelin MPF while no effects were observed for individuals without a FH. These risk-effects on apparent myelin were moderated by hypertension and inflammation-related blood markers (Mole et al., 2020).

The precise nature of these complex synergistic effects between obesity and *APOE* ε4 remain elusive and require further investigation. A recent animal study (Jones et al., 2021) points to inflammation and neuronal plasticity mechanisms underpinning interaction effects between obesity and *APOE* genotype. In this study, a high-fat diet increased gliosis and immediate-early gene expression only in *APOE* ε3 but not *APOE* ε4 knock-in mice. This suggested early dysregulation of adaptive inflammatory mechanisms in *APOE* ε4 mice that may make the brain more vulnerable to insults and damage in the long run. In addition, *APOE* ε4 is also known to lead to changes in glucose, insulin, and lipid metabolism and altered beta-amyloid production (Jones and Rebeck, 2018; Jones et al., 2019). Together, these findings suggest that *APOE* ε4 and obesity lead to metabolic alterations, including inflammatory processes, and may adversely interact with other and with other genetic factors on brain structure and function. We propose that interaction effects between *APOE* ε4 and obesity on the hippocampal formation may increase that region’s vulnerability to subsequent neurodegeneration.

The effects of risk on the hippocampal formation were observed while accounting for age and sex and interaction effects between WHR and *APOE* were only present for the size/complexity component. In contrast, aging was associated with a non-linear inverted U-shaped curve of the myelin/neurite packing component akin to the trajectory of white matter microstructure across the lifespan in humans and rhesus monkeys (Bartzokis et al., 2010; Kubicki et al., 2019; Slater et al., 2019; Yeatman et al., 2014). Myelin/neurite packing increased between the 30s and 40s, when it peaked, remained relatively stable in the 50s and 60s and declined in the 70s (Figure 4A). Age effects were present in all four subfields of the hippocampal formation, i.e., CA1, CA2/3, DG, and subiculum. We propose that this observed age trajectory in the myelin/neurite packing component most likely reflects the maturation of white matter pathways within the hippocampal formation, such as the perforant path, mossy fiber, and Schaffer collateral pathways (Zeineh et al., 2017). Indeed, age-related differences in the myelin basic protein (MBP) expression were found in the CA1, CA2/3 and DG regions of gerbils such that white matter fibers of the perforant pathway, the mossy fibers and Schaffer collaterals were reduced in older relative to younger gerbils (Ahn et al., 2017) Our findings accord with a study (Douaud et al., 2014) that employed a data-driven analysis of gray matter structural variation and identified a brain network comprising prefrontal, intraparietal, posterior cingulate, and medial temporal lobe regions whose lifespan pattern mirrored brain development and age-related degeneration. The hippocampus forms part of this network which matures during adolescence and young adulthood into midlife and shows heightened vulnerability to accelerated neurodegeneration in older age (Douaud et al. 2014). As we did not observe any age-related differences in the size/complexity and metabolic principal components, we propose that hippocampal changes across midlife and early older age may be primarily driven by changes in white matter myelin and neurite packing rather than a loss of neurons and synapses.

Finally, we tested for the effects of risk factors on cognitive performance and explored the relationship between cognitive performance and hippocampal subfield macro and microstructure. The dimensionality of the cognitive data was reduced to four components reflecting verbal and spatial episodic memory as well as spatial navigation first move latencies and path length. Macro- and microstructural measurements were average over protocol and hemisphere for each subfield. We did not observe any risk effects on cognition. This may suggest that risk-related macro- and microstructural tissue changes in the hippocampal formation precede any cognitive impairment and/or were too subtle to induce any functional impairments in this sample of cognitively healthy adults.

In contrast, age was associated with larger latencies in the spatial navigation task due to older people taking longer to plan and initiate their first move in the virtual water maze. In addition, females performed better in the verbal recall test than males. Finally, we observed significant contributions from CA1 volume and free water content (ISOF) and CA2/3 neurite density (ICSF) to the variance in visual recall component. That meant that individuals with larger volume and neurite density in the CA fields were performing better at visual recall, which required the mental reconstruction of a complex spatial figure arrangement. Thus, although risk factors had no apparent impact on cognition it may be possible that their predispose an individual for future episodic memory deficits.

In summary, we provide novel evidence for dissociations between age and age-independent risk effects on hippocampal subfield macro- and microstructure. Non-linear age effects in myelin/neurite packing were observed in all subfields of the hippocampal formation. Central obesity was associated with reductions in myelin/neurite packing and size/complexity across all subfields, with APOE genotype modifying the effects of obesity on size/complexity. Age and sex were significantly related to performance differences in spatial navigation and recall but no effects of risk factors on cognition were present. Individual differences in CA1 and CA2/3 macro- and microstructure predicted performance in visual recall. It remains to be determined if the observed risk-related hippocampal macro- and microstructural differences may precede any future cognitive decline.

## Acknowledgements

This research was funded by a Research Fellowship awarded to CM-B from the Alzheimer’s Society and the BRACE Alzheimer’s Charity (grant ref: 208). JPA is supported by the Wellcome Trust (grant 103722/Z14/Z) and BMC and SDV’s contribution was supported by Wellcome Trust Senior Research Fellowship awarded to SDV (212273/Z/18/Z). We would like to thank Jilu Mole, Erika Leonaviciute, Fabrizio Fasano, John Evans, Peter Hobden and Sonya Foley-Bozorgzad for their assistance with MRI data acquisition and processing. We would also like to thank Derek A. Hamilton and Adam Hampshire for providing the virtual Morris water maze and Cambridge Brain Sciences battery tasks, Rosie Dwyer, Samantha Collins, Abbie Stark, and Emma Blenkinsop for their assistance with the collection and scoring of the cognitive and health data, and Rhodri Thomas for assistance with the *APOE* genotyping of the saliva samples.

## Conflict of interest

The authors declare no competing financial and/or non-financial interests.

## Notes

### Competing Interest Statement

The authors have declared no competing interest.

